# Electrochemical measurements reveal reactive oxygen species in stress granules

**DOI:** 10.1101/2021.03.16.435640

**Authors:** Keke Hu, Emily Relton, Nicolas Locker, Nhu T. N. Phan, Andrew G. Ewing

## Abstract

Stress granules (SGs) are membraneless organelles that assemble in the cytoplasm to organize cellular contents and promote rapid adaptation during stress. To understand how SGs contribute to physiological functions, we used electrochemical measurements to detect electroactive species in SGs. With amperometry, we discovered that reactive oxygen species (ROS) are encapsulated inside arsenite-induced SGs, and H_2_O_2_ is the main species. The release kinetics of H_2_O_2_ from single SGs and the number of H_2_O_2_ molecules were quantified. The discovery that SGs contain ROS implicates them as communicators of the cellular stresses rather than a simple end-point. This may explain how SGs regulate cellular metabolism and stress responses. This may also help better understand their cytoprotective functions in pathological conditions associated with SGs such as neurodegenerative diseases, cancers and viral infections.

---

Controlling the localization of macromolecules is key for cellular functions and typically achieved by surrounding them with lipid membranes in organelles such as the nucleus. Membraneless organelles such as stress granules (SGs) are increasingly recognized as an alternative way to organize cellular components.^1^ Their formation generates high local concentrations of RNA and protein, providing an ideal platform for the regulation of fundamental processes allowing cells to rapidly adjust in response to various physiological and pathological triggers.^2^

SGs assemble to capture mRNAs and proteins during stresses including oxidative stress, heat shock, viral infection, proteasomal inhibition, ER (endoplasmic reticulum) stress, UV irradiation, among others.^3^ The general inhibition of protein synthesis following stress results in the dissociation of mRNAs from polysomes and their accumulation in ribonucleoprotein (RNP) complexes.^3^ This increases the concentration of cytoplasmic RNPs and their binding by aggregation of prone RNA-binding proteins (RBPs), such as G3BP1 or TIA-1, results in clustering/fusion events driven by interactions between their protein and RNA components, ultimately promoting liquid-liquid phase separation (LLPS) and SG assemblko–y.^4^ SGs are highly dynamic, rapidly dissolving upon stress resolution to release sequestered mRNAs for future translation.

By condensing specific proteins, SGs also alter the composition and concentration of cytoplasmic proteins, affecting the course of biochemical reactions and signalling cascades.^5^ Many signalling molecules associated with diseases concentrate in SGs, suggestive of a role for SGs as signalling hubs.^6^ Furthermore, metabolic enzymes stored in SGs produce metabolites that can affect SG stability such as AdoMet or acetyl-CoA.^7,8^

By concentrating key signalling and cytoplasmic sensors or effectors of innate immunity SGs are at a crossroads between intracellular signalling, antiviral responses and translation control.^9^ Moreover, the dysregulation of SGs assembly is increasingly associated with neuropathologies such as amyotrophic lateral sclerosis (ALS), Parkinson’s and Alzheimer’s diseases.^10–12^ Finally, many SG proteins are aberrantly expressed in tumours and SGs are exploited by cancer cells to adapt to the adverse conditions of the tumour microenvironment.^13^ Therefore SGs are important for many normal and pathological processes. However, beyond their protein and RNA composition, very little is known about the contents of SGs, and how these impact on their functions. With clear functional connections to signalling and metabolism, we speculated that, despite the absence of membranes, SGs could act as stores for metabolites or secondary messengers.

Reactive oxygen species (ROS) are key messengers that trigger and regulate cellular signaling pathways important for a wide range of cellular processes, including proliferation and survival, antioxidant regulation, mitochondrial oxidative stress, apoptosis, aging, iron homeostasis and DNA damage response.^14,15^They are produced by enzymes localized in the cytoplasm, mitochondria, peroxisome, and ER. ROS are a family of molecules that include super oxide radical O_2_·^-^, hydroxyl radical OH· and the freely diffusible H_2_O_2_. H_2_O_2_ is the most stable ROS and regarded as the primary example.^16^ The opposing effects of ROS in physiological processes have attracted considerable attention. They have diverse functions depending on the subcellular resource, location, duration and concentration of these molecules.^16–19^ Electrochemistry offers a direct and simple way to detect ROS as some are electroactive. Normally electro-oxidation is preferred as electro-reduction is susceptible to the interference of oxygen in aerobic environments. ROS detection at single-cell and subcellular levels has been successfully achieved with the development of small probes and new methodologies.^20,21^

Given our limited understanding of SG contents beyond proteins and RNAs, we hypothesized they might contain electroactive species. To test this, we treated human bone osteosarcoma epithelial (U2OS) cells with 500 µM sodium arsenite for 30 min and then isolated SGs as previously described.^22,23^ Electrochemical measurements were carried out with both Pt microelectrodes and platinized carbon fiber microelectrodes (CFME) by either electrochemical collision (“adding”) or direct absorption (“dipping”).^24^ These measurements revealed the apparent presence of ROS within SGs.

The isolation protocol of SGs is shown in the SI. The “granule enriched fraction” including primarily SGs and other proteins was dispersed in homogenizing buffer. A schematic of electrochemical measurement of isolated SGs is illustrated in Figure 1A. An applied potential of 400mV (vs Ag/AgCl) was used as a higher potential might bring more interferences and complicate the measurement. As can be seen from the representative amperometric traces of 33-µm CFMEs and 25-µm Pt electrodes in Figure 1B, current transients were observed only with the Pt electrode indicating that the detected species can be oxidized on the surface of Pt instead of CFME at 400 mV. These species appear to be ROS after excluding other electroactive species such as ascorbic acid, some neurotransmitters, and enzymes.

**Figure 1.**
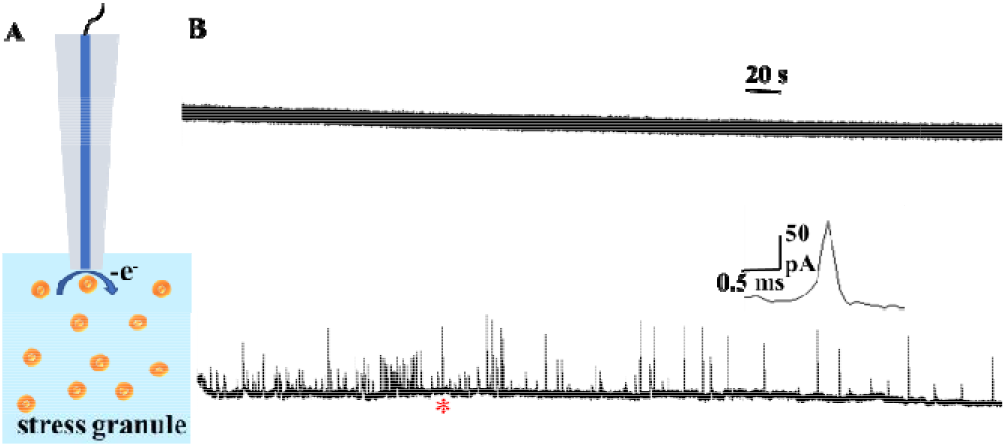
(A) Schematic of electrochemical measurement of stress granules (SGs) isolated from NaAsO_2_-treated U2OS cells; (B) Representative amperometric traces of SGs obtained from a 33µm CFME (upper trace) and a 25µm Pt electrode at 400mV vs Ag/ gCl (lower trace).

Prior to confirming the presence of ROS, we verified that these electroactive species resulted from SGs, rather than impurities or protein contaminants. To this end, SGs were further purified from the SGs-enriched fraction using immunoprecipitation (Figure S1), resulting in pure SG fractions.^22,23^ The schematic of immunoprecipitation is presented in Figure S1. The results shown in Table S1 suggest that the interference from the impurities is negligibly small, and the electroactive species are from the SGs. The Pt microelectrode was replaced with a platinized CFME in order to increase the number of active sites on the electrode surface for ROS oxidation. Representative amperometric traces of Pt black deposition and voltammograms in 10 µM H_2_O_2_ before and after modification are shown in Figure S2. Platinized CFME was then dipped into the suspension of SGs for amperometric measurement. When catalase (CAT, 2.5 mg/mL^25^), the enzyme that converts H_2_O_2_ to O_2_ and H_2_O is added to the solution of SGs, the amperometric response of the same electrode is eliminated providing strong evidence that H_2_O_2_ is the main electroactive species in SGs. As shown in Figure 2A, spikes were observed only in the amperometric trace obtained from the suspension of SGs without CAT.

**Figure 2.**
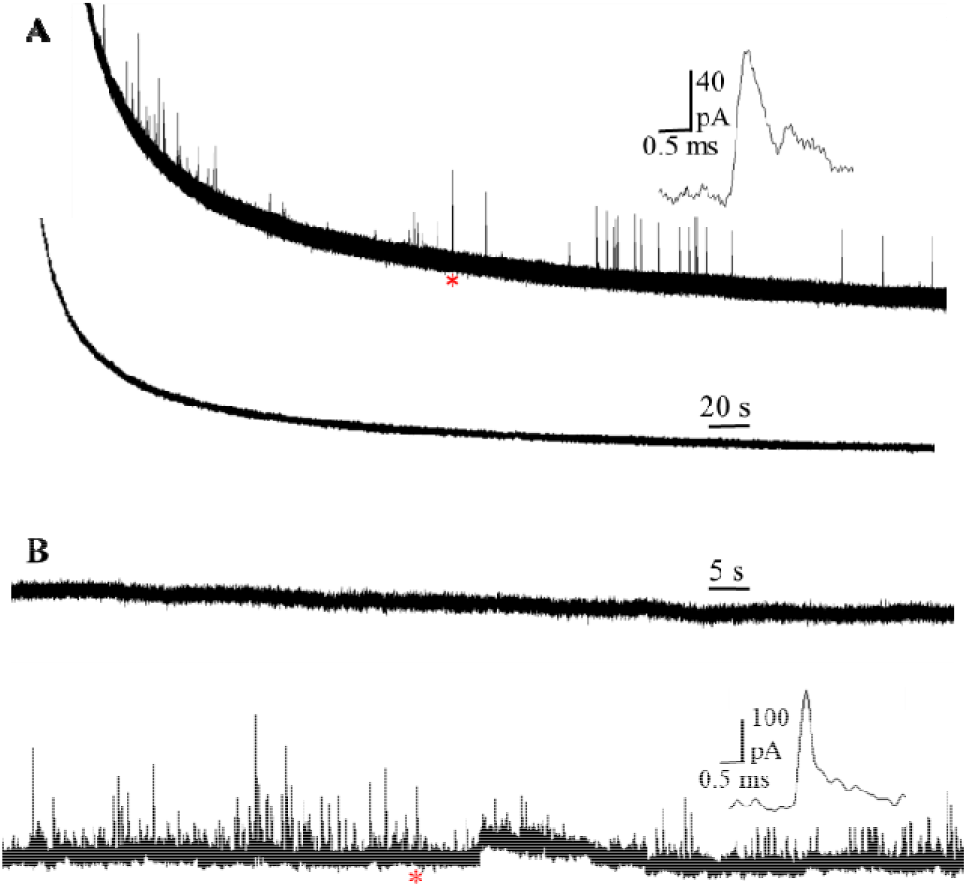
(A) Representative amperometric traces of the same platinized CFME obtained sequentially from SGs suspensions without (upper) and with CAT (lower) at 400 mV vs Ag/AgCl; (B) Representative amperometric traces of another platinized CFME obtained from the two suspensions in a reverse order at 400mV vs Ag/AgCl. The inset shows amplification of the spikes labeled with the red asterisk.

To verify that the disappearance of spikes cannot be attributed to the surface passivation of platinized CFME, another electrode was dipped into the two suspensions in a reverse order. Between these two measurements, the electrode was immersed in PBS solution for 3 min to remove the residues of CAT on the surface. The resultant amperometric traces are presented in Figure 2B with spikes only from SGs suspension without CAT (lower trace). It should be noted that a small number of spikes were captured in some traces obtained from SGs suspension with CAT (Figure S3), indicating that in addition to H_2_O_2_, there might be small amount of other species in SGs.

However, we conclude that the main electroactive species in arsenite-induced SGs is H_2_O_2_.

To quantify the release kinetics of H_2_O_2_ from single SGs and the number of H_2_O_2_ molecules, spikes with a single peak were analyzed. A typical spike analysis is illustrated in Figure 3A with I_max_ of 22.8 pA, *N*_molecules_ of 3.82 10^4^ and release kinetics of 457.8 µs. The shape of the transient matches well with theoretical curves (the red one). Statistical analysis of 229 release events shows the median for t_1/2_ is 0.48 ms. The distribution of log [*N*_molecules_] is presented in Figure 3B with a Gaussian distribution. The *N*_molecules_ ranges from 1×10^4^ to 1.2×10^5^ with the median of 4.03×10^4^.

**Figure 3.**
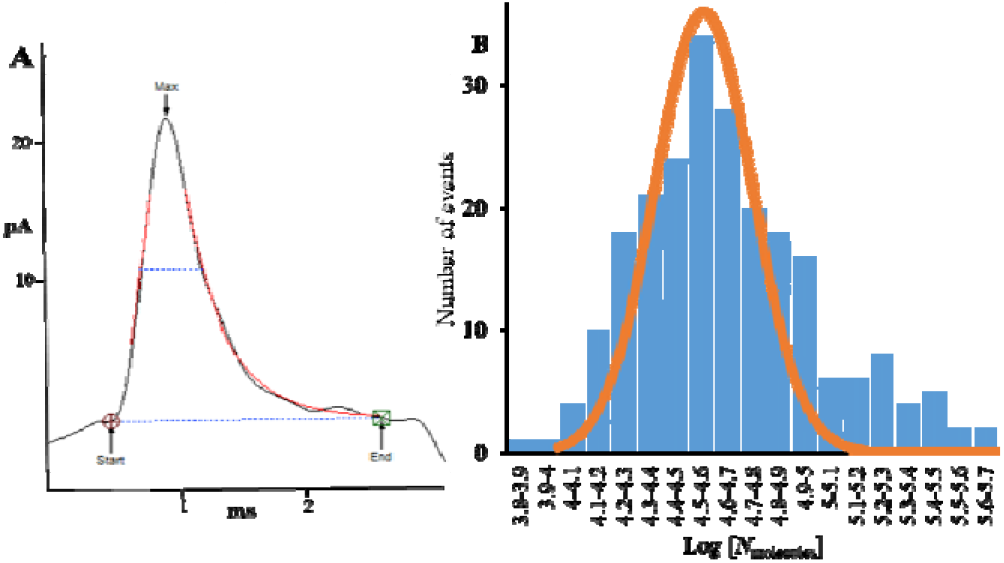
(A) A typical spike analysis for quantification of release kinetics of H_2_O_2_ from single SGs and the number of H_2_O_2_ molecules; (B) Distribution of log [*N*_molecules_]. Fits were obtained from a Gaussian distribution of the data. (collected from four isola ions of SGs; number of events, 229).

Because cancer cells are typically enriched with ROS,

we next assessed whether SGs assembled in non-cancer cells also contain ROS. To test this, we used MCF-10A non-cancer cells, which, unlike cancer cells, contain no ROS or the amount is too low to be detectable without any treatment.^26^ First, we established that arsenite stimulation of MCF-10A cells resulted in SG assembly, as detected by the accumulation of cytoplasmic G3BP1 foci (Figure 4A). Next, we performed amperometric analysis of the isolated SGs. Following stimulation with arsenite and SG assembly, similar results were obtained from SGs formed inside these non-cancer cells with spikes appearing only from platinized CFME instead of C ME (Figure 4B). Thus, this demonstrates that both cancer and non-cancer cell lines form SG that contain ROS upon stimulation with arsenite.

**Figure 4.**
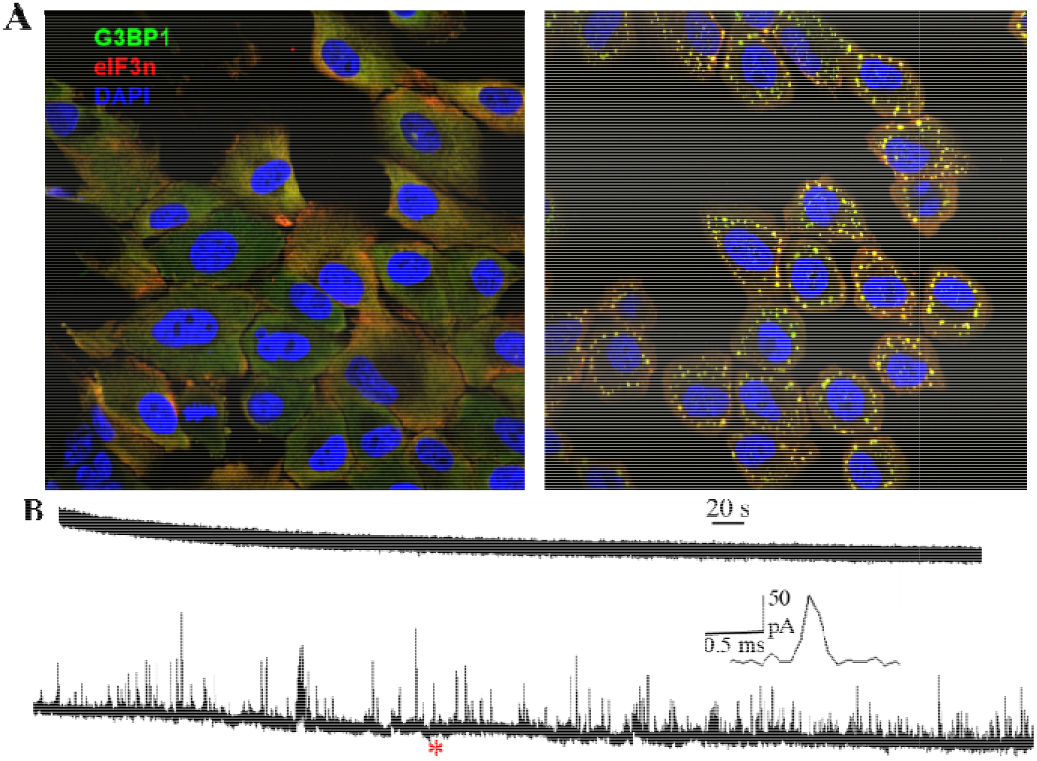
(A) Representative fluorescence microscopy images of MCF-10A cells without (left) and with (right) NaAsO_2_ treatment (yellow dots represent SGs). (B) Representative amperometric traces of SGs isolated from a non-cancer cell (MCF-10A) obtained from CFME (upper) and platinized CFME at 400mV vs Ag/AgCl (lower), the inset shows an amplific tion of the spike labeled with the red asterisk.

Both SGs and ROS play pivotal roles in redox regulation and cell signaling. Many signaling molecules associated with diseases can concentrate in SGs, with several shown to be regulated by SGs. The accumulation of the pro-apoptotic RACK1 results in the supression of MTK1-sPAK signaling and a blocking of the resulting apoptosis.^27^ Similarly, heat shock results in the assembly of SGs that trap TRAF2, impairing NF-ĸB dependent proapoptotic proinflammatory responses.^28^ Proinflammatory and responses are also silenced by the sequestration within SGs of SRC-3, a regulator of several transcription factors promoting inflammation.^29^ These studies support a role for SGs as signaling hubs, and they are generally considered cytoprotective.^6,27,30^

H_2_O_2_ has been identified as a second messenger molecule in subcellular compartments.^31^ Nicotinamide adenine dinucleotide phosphate (NADPH) oxidase (Nox) is responsible for the generation of ROS in immune cells during phagocytosis. However, nonphagocytic Nox enzymes have been found to be involved in many subcellular locations such as the endoplasmic reticulum, nucleus, and mitochondria.^32–34^ One or more Nox isoforms could be located in several subcellular compartments within a single cell type.^35^ And, expression of Nox enzymes in subcellular domains can regulate their participation in varied signaling pathways. The influence of compartmentalized Nox in immune signaling pathways and the compartmentalization of H_2_O_2_ in survival signaling have been reported.^36–38^ Further studies should aim at exploring potential interactions between Nox enzymes and stress granules.

The localization of ROS in SGs reveals the role of SGs as signaling hubs, and these ROS might participate in SGs-associated physiological activities. While high concentration of H_2_O_2_ triggers SGs assembly, the presence of H_2_O_2_ within SGs could be involved in feedback loop control of SG assembly by oxidizing essential proteins in the SGs, as previously shown for TIA1. This might sensitize the cells to a variety of pathological processes.^39^ Aging neuronal cells and tumor cells are more prone to encounter ER stress or oxidative stress. Consequently, ROS-mediated disturbance of SG formation could impair the cytoprotection provided by SGs and promote degeneration initiated by these SG-inducing as well as SGnon-inducing stresses.

In conclusion, using amperometry we have discovered that arsenite-induced SGs contain ROS in both cancer and non-cancer cells and that H_2_O_2_ is the main electroactive species in SGs. This significantly advances our understanding of SG composition beyond macromolecules such as RNA and proteins. The discovery of these small molecules will help uncover new functions for SGs associated with cellular physiology. These ROS might participate in SGs-related activities including redox regulation and cell signaling. Yet, in some tumor cells and especially aging neuronal cells, ROS might impair SGs-mediated cytoprotection by oxidizing essential components of SGs and render the cells sensitive to pathological insults, which will promote neurodegeneration. Although the exact biological function of SGs remains poorly understood, this finding unveils a correlation between SG biology and pathogenesis in NDs and some cancers, which will offer new insights into the therapeutics of SG-related diseases. As ER stress and oxidative stress are implicated in other diseases such as diabetes, stroke and atherosclerosis, the malfunction of SGs formation might also be involved in these disorders.

Given the indication that the composition of SGs can differ significantly depending on cell type and stress stimuli, further investigation is needed to establish how SG heterogeneity impact on their capacity to store reactive species, and their impact on cellular function.

## ASSOCIATED CONTENT

### Supporting Information

Experimental details, schematic of immunoprecipitation, scheme and result of verification of ROS from SGs, platinization of CFME and voltammograms in H_2_O_2_ solution, additional amperometric trace of platinized CFME in SGs suspension with added CAT including Figures S1-S3 and Table S1. This material is available free of charge via the Internet at http://pubs.acs.org.

## AUTHOR INFORMATION

### Notes

The authors declare no competing financial interests.

## ACKNOWLEDGMENT

We acknowledge funding from the European Research Council (Advanced Grant), the Knut and Alice Wallenberg Foundation, the Swedish Research Council (VR), the National Institutes of Health, and the Medical Research Council (MR/R02426X/1).

## TOC Graphic

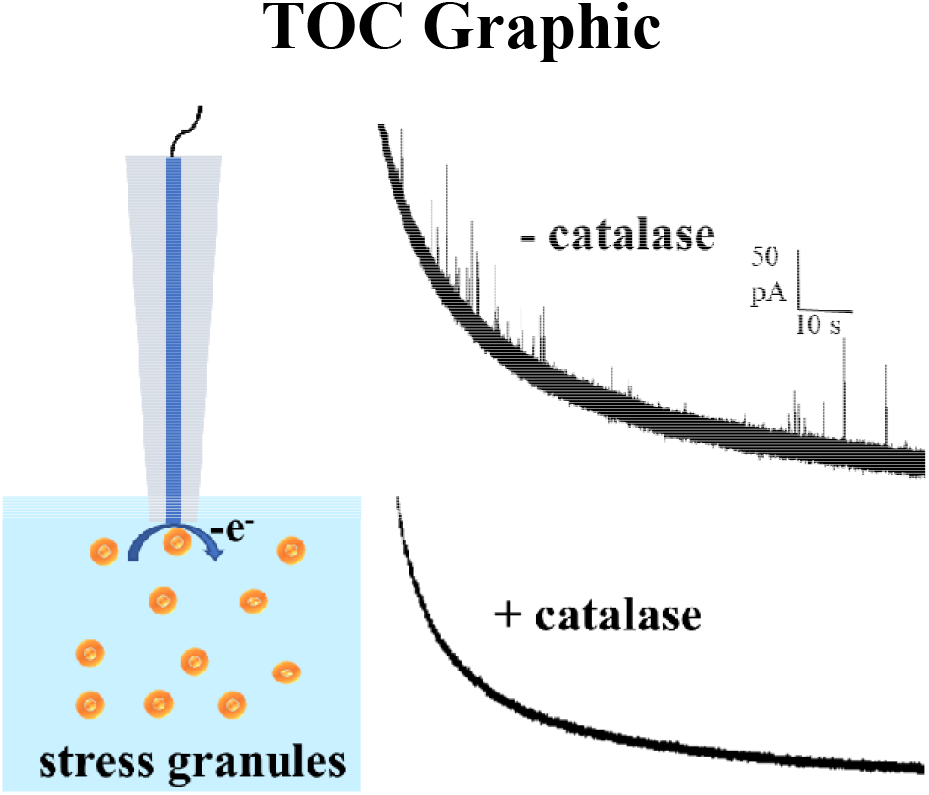

