## Supplementary material for "Electrochemical measurements reveal reactive oxygen species in stress granules"

### Table of contents


1. **Chemicals**
2. **Experimental**

   **2.1 Cell culture**

   **2.2 SGs isolation protocol**

   **2.3 Immunofluorescence microscopy imaging of MCF-10A cells**

   **2.4 Fabrication and platinization of 33-µm CFME**

   **2.5 Data acquisition and analysis**
3. **Verification of ROS from SGs**

   Figure S1. Schematic of immunoprecipitation for SGs purification

   Table S1. Statistical results for amperometric measurements of impurities with Pt electrodes
4. **Fabrication of CFME and testing in $H_2O_2$ solution**

   Figure S2. Platinization of CFME and voltammograms in 10 µM $H_2O_2$
5. **Amperometric trace of platinized CFME in SGs suspension with CAT**

   Figure S3. Representative amperometric trace of platinized CFME in a SGs suspension with CAT added


### 1. Chemicals

All reagents were purchased from Sigma-Aldrich unless otherwise specified. Phosphate buffered saline (PBS, 10 mM $Na_2HPO_4$, 2.7 mM KCl and 137 mM NaCl, pH =7.4) was prepared by dissolving one tablet in 200 mL deionized water (Milli-Q; Millipore Corp.) and used as supporting electrolyte in CFME platinization and in hydrogen peroxide ($H_2O_2$, 30 wt. %) dilution.

Homogenizing buffer (in mM): 230 sucrose, 1 EDTA, 1 $MgSO_4$, 10 HEPES, 10 KCl, cOmplete enzyme inhibitor (Roche, Sweden), DNase I (10 µg/mL) (Roche), 0.001 oligomycin, pH 7.4, ~310 mOsm.

### 2. Experimental

#### 2.1 Cell culture

Human bone osteosarcoma epithelial (U2OS) cells were maintained in high glucose Dulbecco's modified Eagle's medium (DMEM), supplemented with 10% fetal bovine serum (FBS), 1 % penicillin-streptomycin and 1ug/mL puromycin at 37°C in a 5 % $CO_2$ environment (all supplements purchased from Invitrogen).

A human non-tumorigenic epithelial cell line MCF 10A (ATCC® CRL-10317™) was maintained in DMEM, 4,5g/L D-glucose, Na pyruvate, L-glutamine supplemented with 10 % fetal calf serum (FCS), 100 U of penicillin/mL, 100 µg of streptomycin/mL, 10 mM HEPES, and 2 mM L-glutamine (all supplements purchased from Invitrogen) at 37°C in a 5 % $CO_2$ environment.

#### 2.2 SGs isolation protocol [1,2]

U2OS cells were stressed with arsenite (0.1 mM for 1 h) then were crosslinked with 1 % formaldehyde in PBS for 10 min at room temperature (RT), and quenched by adding 125 mM of glycine for 10 min at room temperature.

The cells were spun at 4°C for 5 min at 230 g and the pellet was re-suspended in 1 ml of SG lysis buffer (50 mM Tris-HCl pH 7.4, 100 mM potassium acetate, 2 mM magnesium acetate, 0.5 mM DTT, 50 µg/mL heparin, 0.5 % NP-40, 1 complete mini EDTA-free protease inhibitor tablet (Roche)) and then lysed by passing through a 25 G 5/8 needle seven times on ice and spun at 1,000 g for 5 min. The supernatant was collected and spun down at 18,000 g for 20 min at 4°C. Subsequently the supernatant was removed and the pellet was re-suspended in 1 mL of SG lysis buffer and again spun down at 18,000 g for 20 min at 4°C. Finally, the pellet was re-suspended in 300 µL of SG lysis buffer (stress granule core enriched fraction) and nutated at 4°C for 1 h with 60 µL of magnetic Dynabeads Protein A (Invitrogen). Once the beads have been removed, the supernatant was incubated with specific antibody (1 µg of rabbit anti-GFP antibody and 1 µg of IgG, Invitrogen, Monoclonal mAb 3E6) and nutated overnight at 4°C. The unbounded antibody was removed by centrifugation at 18,000 g for 20 min at 4°C, and the pellet was re-suspended in 300 µL of SG lysis buffer before to incubate it with 60 µL of Dynabeads Protein A at 4°C for 3 h. The Dynabeads were then washed for 2 min at 4°C with 1 mL of wash buffer 1 (SG lysis buffer and 2M Urea), for 5 min at 4°C with 1 mL wash buffer 2 (SG lysis buffer and 300 mM potassium acetate), for 5 min at 4°C with 1 mL SG lysis buffer and seven times with 1 mL of TE buffer (10 mM Tris HCl pH 8.0, 1 mM EDTA pH 8.0).

#### 2.3 Immunofluorescence microscopy imaging of MCF-10A cells

Approximately $5x10^5$ MCF-10A cells were plated on glass coverslips in a 24 well plate. Following arsenite stimulation, cells were fixed with a solution of 4 % PFA (Sigma) in PBS for 20 min at RT, washed in PBS and stored at 4°C. After permeabilization with 0.1 % Triton-X100 (Sigma) in PBS for 10 min at RT and 3 washes with PBS, the coverslips were blocked with 0.5 % BSA (Fisher) in PBS for 1 h at RT then incubated with 200 µL of primary antibodies solution for 1 h at RT. After 3 washes with PBS, coverslips were incubated in the dark with 200 µL of fluorescently-labelled secondary antibodies solution containing 0.1 µg/mL DAPI solution (Sigma). The coverslips were then washed and mounted on slides with a drop of Mowiol 488 (Sigma). Confocal microscopy was performed on a Ti-Eclipse - A1MP Multiphoton Confocal Microscope (Nikon) using the Nikon acquisition software NIS-Elements AR. MCF10A cells, with or without sodium arsenite treatment, were fixed and stained with primary antibodies specific for G3BP (rabbit anti-G3BP, Invitrogen) and eIF3B (goat anti-eIF3B, Santa Cruz). This was followed by staining with Alexa Fluor 488-conjugated donkey anti-rabbit IgG (Invitrogen) and Alexa Fluor 555-conjugated donkey anti-goat IgG secondary antibodies (Invitrogen). Nuclei were stained with DAPI.

#### 2.4 Fabrication and platinization of 33-µm CFME

CFME Fabrication

A 33-µm diameter carbon fiber was aspirated into a borosilicate capillary (1.2 mm O.D., 0.69 mm I.D., Sutter Instrument Co.). The capillaries were subsequently pulled in half with a micropipette puller. The electrode was then sealed by dipping the pulled tip in epoxy (Epoxy Technology) solution. The glued electrodes were cured in an oven at 100°C overnight and subsequently cut at the glass junction and beveled at 45° angle (EG-400, Narishige Inc.).[3]

Platinization of CFME

The stock solution contained 125 µL hexachloroplatinic acid (8 wt. %; Sigma-Aldrich) and 0.2 mg lead (II) acetate trihydrate (99.8 %; Sigma-Aldrich) in 6.4 mL PBS. It was diluted 5 times with PBS before platinization. The polished CFME was biased to -130 ± 70 mV vs. Ag/AgCl reference using CHI electrochemical analyzer (CH Instruments, Inc.). The extent of platinization was controlled by monitoring the reductive current, and the process was interrupted when the electrical charge of the signal reached the desired value of ~300 µC.[4] Platinized microelectrodes were stored in PBS before use.

#### 2.5 Data acquisition and analysis

Current transients were recorded and digitized using a Digidata1440A (Molecular Devices) and digitized at 10 kHz or 100 kHz and filtered at 2 kHz using a 4-pole Bessel filter. The data were converted in Matlab (The MathWorks, Inc.) and processed in IgorPro (Wavemetrics, Lake Oswego, OR).[5] Traces were manually checked for potential false detections done by the software. Spike characteristics were determined as number of molecules based on the charge measured in each spike, $t_{1/2}$ = full spike width at half-maximum. The medians were calculated from the current transients of all SGs analyzed.

### 3. Verification of ROS from SGs

SGs were purified from the SGs-enriched fraction using immunoprecipitation (described above), resulting in pure SG fractions. The schematic of immunoprecipitation is presented in Figure S1. After purification was done, the impurities were taken for amperometric measurement with Pt microelectrodes. As shown in Table S1, for more than half of the electrodes, no spikes were observed, and only one electrode gave 3 spikes. This suggests that the interference from the impurities is negligibly small, and the electroactive species are from the SGs.

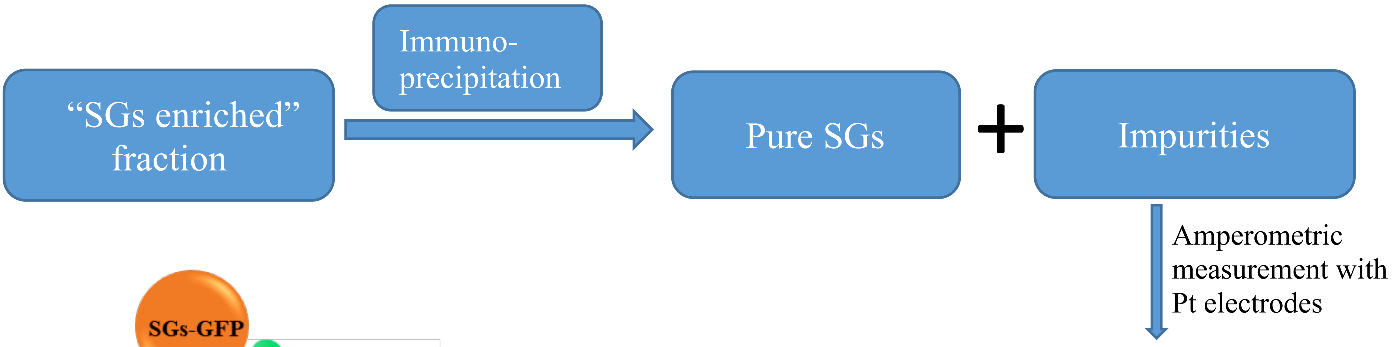


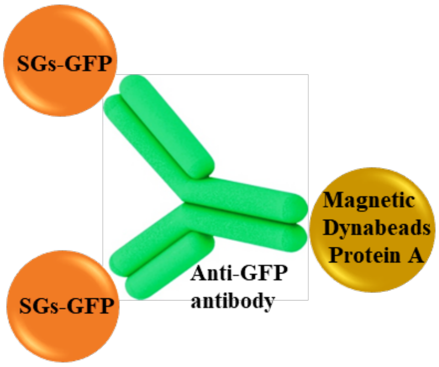


Figure S1. Schematic of immunoprecipitation for SGs purification

| | | | | |
|---|---|---|---|---|
| **Number of electrodes** | 19 | 10 | 4 | 1 |
| **Number of spikes** | 0 | 1 | 2 | 3 |

Table S1. Statistical results for amperometric measurements of impurities with Pt electrodes (collected from 4 isolations of SGs, 34 Pt electrodes)

### 4. Platinization of CFME and testing in $H_2O_2$ solution

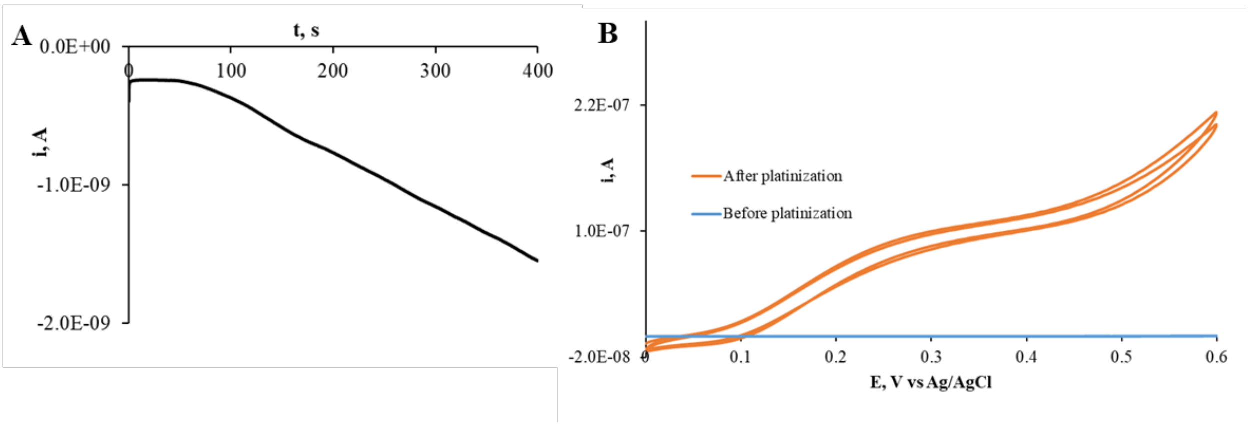


Figure S2. (A) Representative amperometric trace for a 33-µm CFME platinization. (B) Representative voltammograms of CFME in 10 µM $H_2O_2$ solution before (no response) and after platinization

### 5. Amperometric trace of platinized CFME in a SGs suspension with CAT

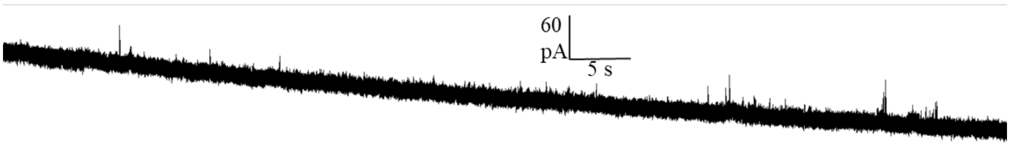


Figure S3. Representative amperometric trace of platinized CFME in a SGs suspension with CAT added.
